# NGSpop: A desktop software that supports population studies by identifying sequence variations from next-generation sequencing data

**DOI:** 10.1101/2021.11.22.469506

**Authors:** Dong-Jun Lee, Taesoo Kwon, Hye-Jin Lee, Yun-Ho Oh, Jin-Hyun Kim, Tae-Ho Lee

## Abstract

Next-generation sequencing (NGS) is widely used in all areas of genetic research, such as for genetic disease diagnosis and breeding, and it can produce massive amounts of data. The identification of sequence variants is an important step when processing large NGS datasets; however, currently, the process is complicated, repetitive, and requires concentration, which can be taxing on the researcher. Therefore, to support researchers who are not familiar with bioinformatics in identifying sequence variations regularly from large datasets, we have developed a fully automated desktop software, NGSpop. NGSpop includes functionalities for all the variant calling and visualization procedures used when processing NGS data, such as quality control, mapping, filtering details, and variant calling. In the variant calling step, the user can select the GATK or DeepVariant algorithm for variant calling. These algorithms can be executed using pre-set pipelines and options or customized with the user-specified options. NGSpop is implemented using JavaFX (version 1.8) and can thus be run on Unix like operating systems such as Ubuntu Linux (version 16.04, 18.0.4). Although there are several pipelines and visualization tools available for NGS data analysis, most integrated environments do not support batch processes; thus, variant detection cannot be automated for population-level studies. The NGSpop software, developed in this study, has an easy-to-use interface and helps in rapid analysis of multiple NGS data from population studies.

## Introduction

Next-generation sequencing (NGS) is widely used in all areas of genetic research, such disease diagnosis and breeding, this is in part because it is a useful tool for the detection of sequence variations [1-3]. NGS technology was originally used to study individuals and small samples, but more recently, it has been used to study cohort-level populations. In a medical study, such as that by the Undiagnosed Diseases Network (UDN) showed that a genetic diagnosis with NGS is valid, even if the disease is undiagnosed [4]. According to NGS, 21% were changed in therapy, 37% in diagnostic testing, and 36% in variant-specific genetic counseling. NGS has also been used to construct an ultra-high-density genetic map for the identification of molecular markers for agricultural research [5,6]. The research showed that a genetic breeding with NGS is a valid and reliable tool to develop useful characters. NGS produces a large amount of data, especially for studies involving genetic diseases and breeding at the population level. The identification of sequence variants in these large datasets is one of the most important processing steps; however, currently, sequence variation detection is both complicated and repetitive. Genomics consortia, such as the 1000 genome project [7], provide shell scripts that implement a standard operation procedure (SOP) for variant detection, which helps to standardize the process (https://github.com/ekg/1000G-integration). However, most of the SOP shell scripts in use are difficult to understand and automate. There are several workflows and tools available that include quality control (QC), mapping and the calling, annotation, and visualization of variations. Some tools have too many functions, and consequently, they can be difficult to learn and often require official training. Furthermore, for some tools, the lack of tool integration, and the many options included in their functionality, can confuse the user and considering the available options can be time consuming. Many pipelines and workflows have been developed by commercial and open-source communities to support NGS data analysis. Pipelines such as the ngs_backbone [8] and GATK [9] provide simple commands to perform a complete NGS data analysis. Most pipelines offer only a command-line interface, and thus the user needs to be trained in Unix/Linux commands, shell scripts, or Python. It is difficult to automate variant detection in population-level studies. Galaxy [10] and the CLC genomics workbench [11] provide users with easy-to-use graphical user interfaces (GUIs). Although there are many pipelines and integrated environments for NGS data analysis, each has its own strengths and limitations (Table 1).

**Table 1.** Comparison of the user-friendly graphic interfaces and functions of the SNP analysis pipelines.

| Analysis \ Name | Annovar | Ngs_backbone | inGAP | Galaxy | CLC<br>genomics<br>workbench | NGSpop |
| --- | --- | --- | --- | --- | --- | --- |
| Quality Control (QC) |  | O | O | O | O | O |
| Read Mapping |  | O | O | O | O | O |
| Variant calling<br>(GATK) |  | O | O | O | O | O |
| Variant calling<br>(DeepVariant) |  |  |  |  |  | O |
| Variant annotation | O |  |  | O | O | O |
| Visualization |  |  |  |  |  | O |
| Manual mode | O | O | O |  | O | O |
| Batch mode |  |  |  | O | O | O |

To support sequence variation detection in population-level genomics studies, we have developed a desktop software, NGSpop. The software accepts multiple NGS datasets and allows the user to select between the GATK or DeepVariant [12] calling algorithms. The functionalities for variant detection include QC, mapping, filtering, variant calling, and visualization. Moreover, NGSpop has two modes of action: a one-step mode that supports batch identification of variants and a step-by-step mode in which the user can verify the result of each step. When the user selects the one-step mode, NGSpop can be executed using pre-set options to exclude the time-consuming steps. NGSpop can only be used with Linux operating systems.

### Implementation

NGSpop was implemented using JavaFX (version 1.8), and the tools employed within it were compiled on Ubuntu Linux (version 18.0.4). The GNU compiler collection version 7.2.0, for Ubuntu Linux, was used as a C-language compiler.

### Tools used in the pipeline

The tools included in NGSpop were carefully chosen according to the pipeline of the National Agricultural Biotechnology Information Center (NABIC, Republic of Korea; Fig. 1). NGS data need to be evaluated for QC, and for this purpose, NGSpop includes FastQC (version 0.11.5). Filtering and trimming of the NGS data is mandatory, depending on the sequence quality, and for this step, NGSpop employs TrimmOmatic (version 0.36) [13]. After the QC step, sequence reads can be mapped in NGSpop against a reference genome using an alignment tool, such as BWA (version 0.7.16a) [14], and SAMtools [15] is used for file format conversion and indexing. Mate-pair information cannot be concordant with the sample library information and should be fixed. If sequence reads can be mapped to more than two loci, then the duplicate reads should be removed, and Picard (version 2.9.4) is used for this in NGSpop. For SNP/INDEL identification, the user can select SNP/INDEL identification algorithms from the Genome Analysis Toolkit (version 3.7.0) or DeepVariant (version 0.5.1). Currently, DeepVariant is only supported by the Linux operating system, and consequently, this system is required to run NGSpop. To annotate the identified SNP/INDELs, SnpEff is used (version 4.3q) [16]. The identified and annotated variants are visualized using JBrowser software (version 1.12.3) [17]. All the tools integrated into NGSpop are summarized in Table 2.

**Fig 1.**
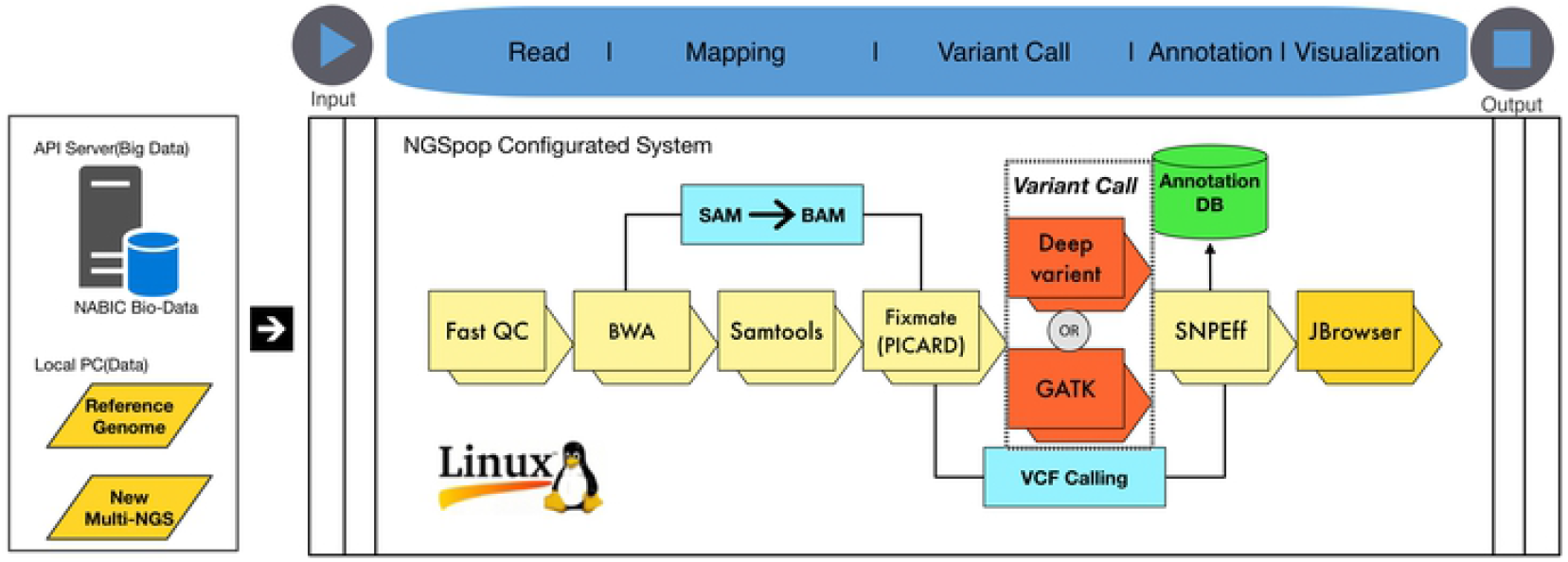
NGS data analysis pipeline used in the NGSpop software. The variant analysis protocol and tools are chosen according to the pipeline of the National Agricultural Biotechnology Information Center (NABIC, Republic of Korea).

**Table 2.**
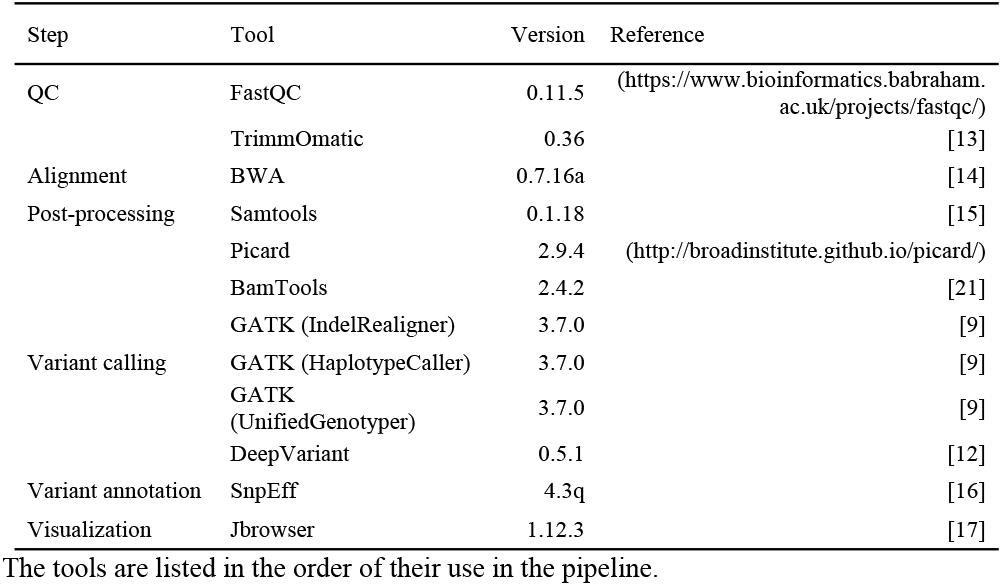
Tools included in the NGSpop software.

### Project creation and importing input files

The user must create a project and specify the data files, including fastq files of sequencing reads and a reference file in the FASTA format (Suppl. 2a). Fastq files of the sequencing reads can be multiple pairs of forward and reverse reads for population studies. Only fastq files produced by the Illumina platform can be processed using NGSpop. For the convenience of users, NGSpop can download a reference file from a genomic database such as the NCBI, Ensemble, and the NABIC server through the application program interface. When the index file of the reference sequence does not exist, NGSpop performs indexing of the reference file.

### Step-by-step mode

NGSpop provides the user with a step-by-step mode, in which they can investigate each step of the analysis. The user can change or execute each option during each step, and changes will become the default options for the same step in each subsequent run (Suppl. 2b). To monitor the progress of each step, NGSpop provides the user with a log window.

### One-step mode

To automate NGS data analysis and support the largescale identification of variants, NGSpop provides a one-step user mode that can run all processes employed by NGSpop with a single click. When NGSpop runs using the one-step mode, the default options will be used for each step. The user can customize the default options used in the one-step mode by first using the step-by-step mode.

### Quality control

To identify highly accurate genomic variation information from the population, the quality of the NGS data should be carefully checked and filtered; FastQC (version 0.11.5) is used for this purpose in NGSpop. Sequence reads that are below the score (Phred) [18] specified by the user will be filtered out and low-quality regions at the 5′- and 3′-ends can be trimmed using TrimmOmatic (version 0.36) [13].

### Read mapping and duplicate removal

NGSpop employs BWA (version 0.7.16a) [14] as a mapping tool for the NGS reads. To convert the BWA sequence alignment map format (sam) to a binary alignment map (bam) format, and then to sort and index the file, NGSpop uses SAMtools [15]. If the mate-pair information is not concordant with the sample library information, it should be verified and fixed. For this purpose, the Fixmate command of Picard (version 2.9.4) is used in NGSpop. In addition, duplicate reads are removed using the MarkDuplicates and AddOrReplaceReadGroups commands of Picard, and to calculate the statistics of the sequence reads, BamTools [19] is used.

### SNP/INDEL identification

Using NGSpop, the user can select a SNP/INDEL identification algorithm from the Genome Analysis Toolkit (version 3.7.0) or DeepVariant (version 0.5.1). The Genome Analysis Toolkit (version 3.7.0) [9] is a standard tool for single nucleotide polymorphism (SNP)/INDEL identification from NGS data. To realign the reads around the INDELs, NGSpop, uses the RealignerTargetCreator and IndelRealigner commands of GATK. After the realignment of the reads, UnifiedGenotyper is used as a variant caller in NGSpop.

### DeepVariant

DeepVariant is a variant caller developed by Google Inc. The tool showed overwhelming quality and imputation reference performance compared to well-established pipelines such as GATK [20]. NGSpop includes DeepVariant in its pipeline, and the user can select between the GATK or DeepVariant algorithms. DeepVariant is a deep learning-based variant caller that uses aligned reads (in BAM or CRAM format) to produce pileup image tensors, and each tensor is classified using a convolutional neural network, and finally reports the results in a standard VCF or gVCF file. DeepVariant supports germline variant calling in diploid organisms.

### Variant merge

Vcf files that are produced using the GATK or DeepVariant algorithms in the same project will be merged in NGSpop. The vcf-merge script in VCFtools [20] is employed to integrate the vcf files. VCF tools are a program package of perl modules and C++ programs.

### Variant annotation

Variations in the nucleotides can change the amino acids of the genes and thus affect the organism. Therefore, the functional effects of these variants on the genes should be predicted. To annotate the identified variants, NGSpop uses SnpEff (version 4.3q) [16]. In this study, NGSpop only included the *Arabidopsis thaliana* database (TAIR10 genome [22]) for SnpEff. In other studies, the SnpEff database should be included for the appropriate organism if it is available. If there is no available database for non-model organisms, the database should be generated manually.

### Variant visualization

The annotated variant information can be visualized using JBrowser (version 1.12.3; Fig. 2) [17] in NGSpop. There are four feature tracks in the JBrowser window: reference sequence, annotation information of the reference in GFF, mapped reads in bam format, and annotated variants. The tracks can be shown or hidden by clicking the check box of the corresponding feature tracks that the user wants to investigate. The annotated variant file can be downloaded by clicking the VCF file download button on the top right of the JBrowser.

**Fig 2.**
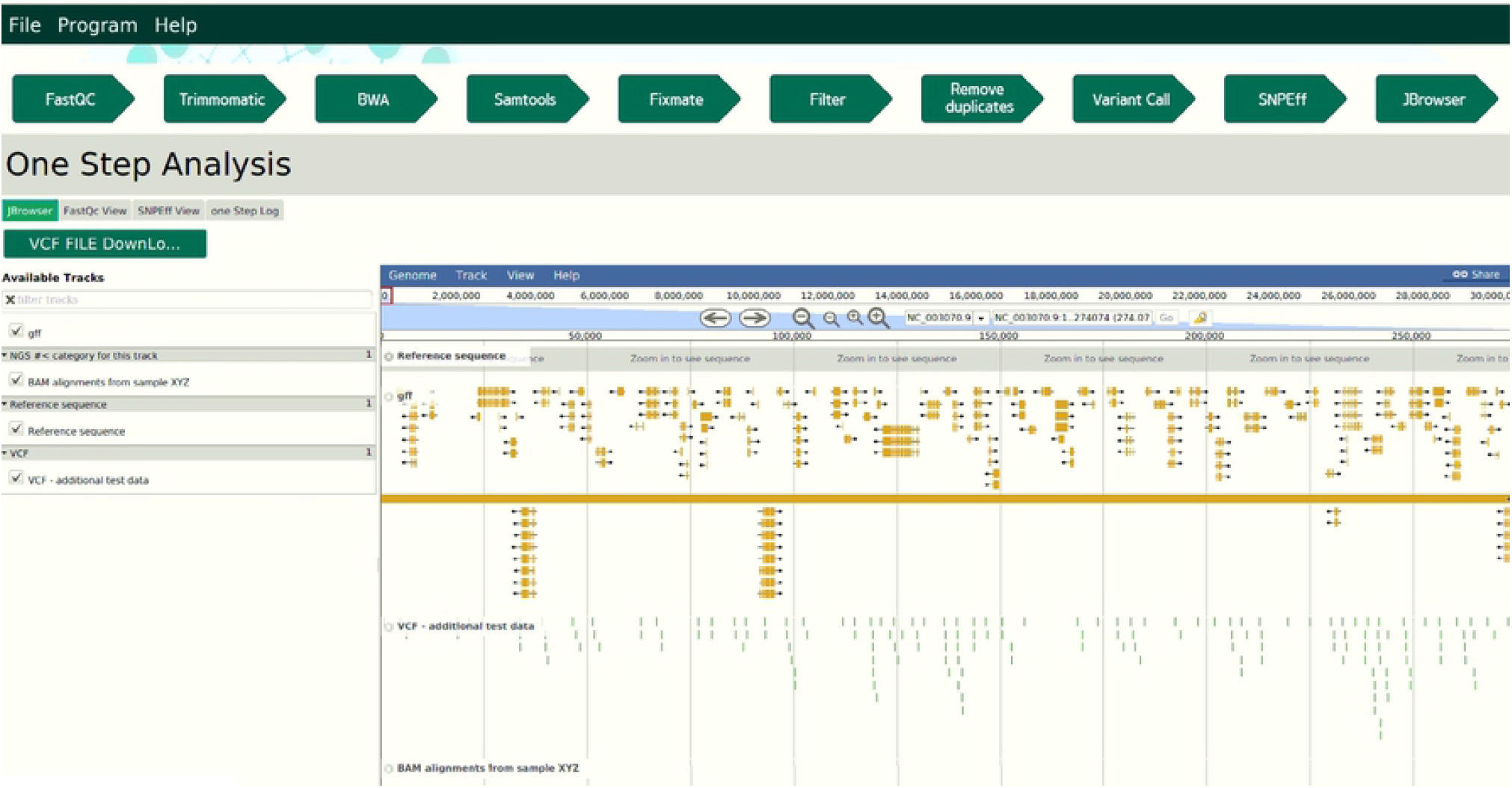
Visualization of variants. Four feature tracks are listed on the left panel of the JBrowser: reference sequence, annotation information of reference in GFF, mapped reads, and annotated variants. Only a .vcf file can be displayed in the JBrowser when multiple NGS data are selected for variant analysis after the merging of multiple .vcf files.

## Results and discussion

The aim of NGSpop is to provide users with an easy-to-use environment for NGS data analysis, regardless of whether the user is an expert in bioinformatics. To this end, NGSpop provides users with two modes: a step-by-step mode for beginners and a one-step mode for experts. There are currently many workflows and easy-to-use tools available for NGS analysis, but as the user is required to run each step manually and wait until each step ends before proceeding, they can slow the rate of analysis. Furthermore, these tools were not designed for population studies, and they only provide users with a step-by-step mode or a difficult hierarchical workflow design. Some tools do provide user-bash script interfaces, but these can be difficult to learn. However, when using NGSpop, only a single click of the run button is required and the results can be visualized using JBrowser. Even though the software provides a user graphic interface, NGSpop accepts multiple pairs of fastq files to support population-level studies. Thereby, users can identify variants in large scale datasets from population studies using only their personal computers (PCs) or workstations. Moreover, NGSpop provides a selection of variant calling algorithms from GATK and DeepVariant in the variant calling steps. Variant calling is an important step in NGS data analysis and genetic studies. There are many tools that identify high-quality and reliable variants from NGS data, but none of them can identify all variants. Therefore, researchers have used and combined multiple tools to identify variants from NGS data. To assess the coverage of the variants identified, we compared the variant calls from GATK and DeepVariant when using NGSpop. For the benchmark test, we generated a total of five test datasets using the complete *Arabidopsis thaliana* genome sequencing data from the DNA Data Bank of Japan (DDBJ) FTP site under the accession number SRR519473 (paired-end run with 52,154,720 reads and 10,430,944,000 bp) [24]. The sequencing data were generated by the *Arabidopsis thaliana* 1001 genome project (http://1001genomes.org) [23] using the Illumina HiSeq 2000 platform. The detailed specifications of the benchmark system and benchmark results are summarized in Table 3. NGSpop took a total of 2 h 46 m 46 s to go from the raw reads to variant annotation or visualization for the five test datasets using GATK, whereas it took 1 h 48 min 00 s when using DeepVariant. A total of 113,163 variants were identified using GATK, and 128,530 variants were identified using DeepVariant. Among the identified variants, 111,918 overlapped between GATK and DeepVariant. Meanwhile, 1,245 and 16,612 were specific to GATK and DeepVariant, respectively. DeepVariant with Tensorflow was faster than GATK in variant calls and identified more variants than GATK. Consequently, NGSpop was found to be an easy-to-use platform for variant calling using GATK and DeepVariant.

**Table 3.** Comparison of the variant calling algorithms used in the NGSpop software.

| Benchmark Machine |  |  |  |  | Variant calling algorithms |  |  |  |  |
| --- | --- | --- | --- | --- | --- | --- | --- | --- | --- |
| CPU | RAM<br>(GBytes) | Storage<br>(TBytes) | OS | Dataset | GATK |  |  | DeepVariant |  |
|  |  |  |  |  | SNP | Time |  | SNP | Time |
|  |  |  |  |  | Count | (hh:mm:ss) | Intersection | Counts | (hh:mm:ss) |
| Intel(R)<br>Xeon(R)<br>CPU E5-2609 v3<br>@<br>1.90GHz | 32 | 1.8 | Ubuntu<br>18.04 | 1 | 20,896 | 0:32:15 | 20,671 | 23,761 | 0:21:19 |
|  |  |  |  | 2 | 20,404 | 0:32:24 | 20,201 | 23,207 | 0:20:04 |
|  |  |  |  | 3 | 22,952 | 0:33:02 | 22,705 | 26,087 | 0:21:38 |
|  |  |  |  | 4 | 24,616 | 0:35:11 | 24,338 | 27,920 | 0:22:35 |
|  |  |  |  | 5 | 24,295 | 0:33:54 | 24,003 | 27,555 | 0:22:24 |
|  |  |  |  | Sum | 113,163 | 2:46:46 | 111,918 | 128,530 | 1:48:00 |
|  |  |  |  | Average | 22,633 | 0:33:21 | 22,384 | 25,706 | 0:21:36 |

Benchmark tests were performed for NGSpop using GATK and DeepVariant as the variant calling algorithms.

## Conclusions

Large-scale parallel sequencing has become a popular tool to identify sequence variations, and many tools have now been developed to analyze NGS data. Although many tools have been developed, few support population-level or cohort-level sequencing data. Owing to the lack of population-level analysis tools, many researchers find it difficult to analyze the massive volumes of NGS data that they produce. Researchers should ideally write scripts to analyze NGS data on the Linux command line. NGSpop is a user-friendly software for researchers who are not familiar with the command line interface and do not want to write shell scripts. Therefore, NGSpop provides the user with an easy-to-use interface and helps to automate the detection of variations from the NGS data at the population level. NGSpop helps genomics researchers who want to analyze population-level NGS data with an easy-to-use GUI. We developed NGSpop to support population-level NGS data analysis; however, there are some limitations. First of all, NGSpop only accepts FASTQ format data that has been produced using Illumina platform because there are too many parameters to consider when analyzing all types of NGS platforms. Next, NGSpop only supports Linux operating system because DeepVariant, one of the variant calling algorithms, can only be used with Linux operating systems. In future studies, we will include functionalities that support NGS platforms other than Illumina while accounting for the variations in formats.

## Availability and requirements

**Project name:** NGSpop

**Project home page:** https://sourceforge.net/projects/ngspop/

**Operating system(s):** Linux

**Programming language:** JavaFX

**Other requirements:** All Perl libraries are listed in the Supplementary information.

**License:** GNU General Public License

**Any restrictions to use by non-academics:** license needed

## Availability of data and materials

**Test datasets**: The test datasets used as the whole-genome shotgun sequencing data are available from the project home page. (https://sourceforge.net/projects/ngspop/).

## Acknowledgements

Not applicable

## Supporting information

**Additional file 1. Supplementary.docx**

## Notes

### Competing Interest Statement

The authors have declared no competing interest.

